# Dynamic distortion of inferred reward probability shapes choice over time

**DOI:** 10.64898/2026.04.08.717213

**Authors:** Matthias Grabenhorst, Laurence T. Maloney

## Abstract

Many choices are triggered by discrete events whose timing determines which options are rewarded. Without informative sensory evidence between events, behavior must rely on internal estimates of latent variables—most notably elapsed time and reward probability. Existing computational frameworks, including evidence-accumulation models, are not designed for this regime, leaving the principles of time-dependent choice unresolved. Here, we formalize choice as an inference problem governed by uncertainty about both elapsed time and reward over time. Participants learned dynamic reward probabilities to guide choices. Behavior approached optimality but exhibited a systematic distortion of inferred reward probability over time, captured by a linear transformation in log-odds space. Crucially, temporal uncertainty was modulated by reward probability but not by elapsed time itself, contradicting Weber-law scaling. These results identify two interacting computational principles–dynamic mapping of reward probability to choice and reward-based temporal precision– that jointly shape behavior when time and reward must be inferred.

## Introduction

Reward expectation is a fundamental determinant of action selection across species and contexts^1-3^. In many natural decisions, however, the probability of reward is not fixed for a given action but varies systematically over time. For example, in animals, the timing of a predator’s sudden attack can determine whether flight or defensive engagement maximizes survival probability. In human social interaction, the timing of a response informs the inferred likelihood of reciprocal interest: a rapid reply invites engagement, whereas a late reply discourages it.

In such settings, reward-maximizing behavior requires resolving two coupled uncertainties: agents must infer where they are in time and what reward structure that temporal position implies. Because elapsed time is internally estimated rather than directly observed^4-7^, reward probability becomes a latent function of another latent variable. Consequently, reward-guided action selection requires joint inference over elapsed time and time-dependent reward probability.

Temporally structured inference of this kind arises across diverse domains, including communication^8^, social coordination^9,10^, skilled motor performance^11,12^, and adaptive interaction with dynamic environments^13^. Across these domains, behavior depends on inferential processes, and, recently, substantial progress has been made in characterizing temporal inference^14,15^, latent-state inference^16,17^, and dynamic representations of expected outcomes^18^, including evidence that neural activity rapidly reflects latent reward structure^19-21^.

However, existing frameworks typically investigate temporal inference and reward inference as separate processes: interval timing models characterize uncertainty in elapsed duration^22,23^, reinforcement-learning, temporal difference models, and related credit-assignment frameworks describe how values are updated across task states^24-27^, and perceptual decision models formalize evidence accumulation under sensory and internal noise^28-31^. In the regime we investigate here, no informative sensory input is available between events, precluding continuous evidence accumulation. Although reinforcement-learning frameworks can, in principle, represent temporal uncertainty through belief states, they are typically not formulated to specify how uncertainty in internal temporal estimates shapes the mapping from dynamic reward probabilities to choice. Thus, it remains unclear how agents combine uncertainty about elapsed time with uncertainty about reward, when reward probability itself changes over time.

Figure 1 formalizes this class of problems: A warning cue establishes a temporal reference and a subsequent target cue occurs at a time point within a bounded interval (Fig. 1a). The timing of this target determines the reward probabilities associated with alternative actions (Fig. 1b, c). To maximize reward, an observer must estimate elapsed time t, map this estimate onto corresponding reward probabilities *P*(*R*|*A*, *t*) and *P*(*R*|*B*, *t*), and select the action with the higher inferred probability. Because reward probabilities vary over time, identical temporal errors can have markedly different consequences depending on when they occur. The core computational problem is therefore not value updating per se, but how temporally indexed, inferred probabilities are transformed into action under temporal uncertainty.

**Fig. 1.**
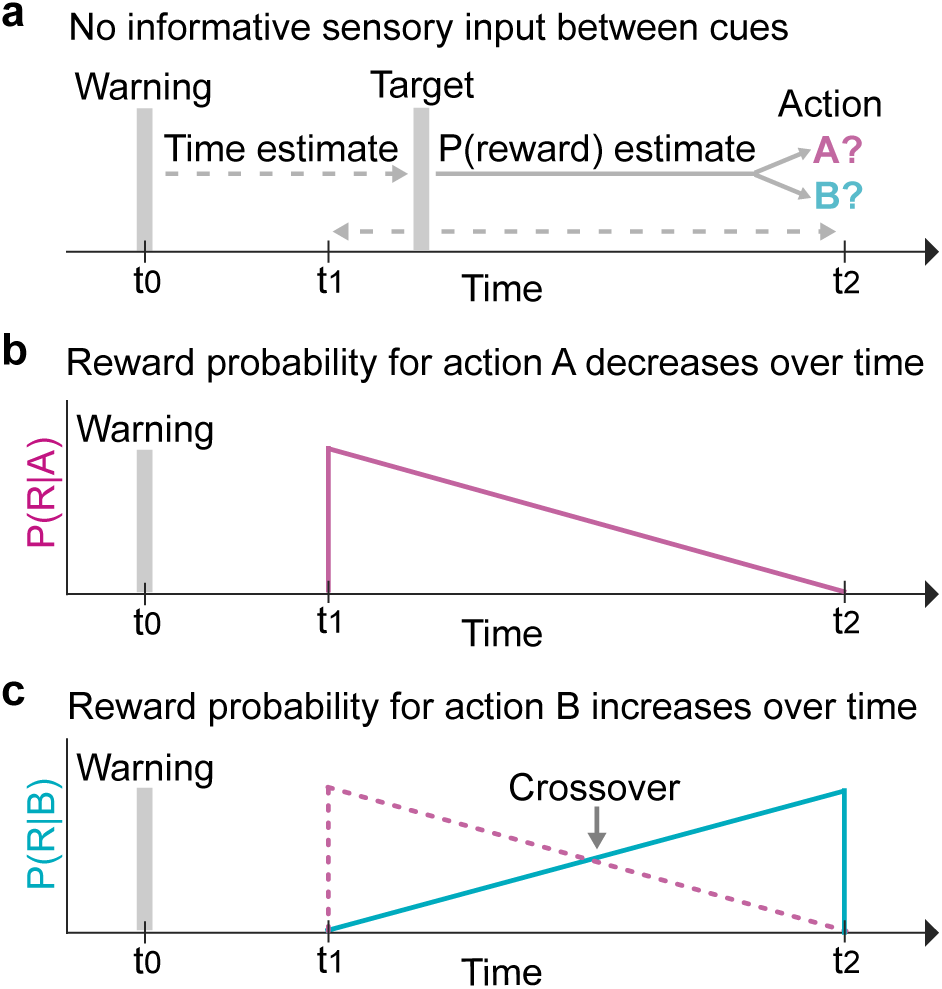
Time-contingent reward inference. **a**, A warning cue defines a temporal reference point (*t*_0_). A target cue occurs at a time *t* within a bounded interval [*t*_1_, *t*_2_]. At target onset, the observer must choose between two actions (A or B). Choice of action therefore relies on internal estimates of elapsed time and inferred reward probability. **b**, The probability of reward associated with action A decreases as a function of elapsed time (illustrated here as linear for clarity). **c**, The probability of reward associated with action B increases over the same interval. At the crossover time point, both actions are equiprobably rewarded. To maximize reward, an optimal observer must estimate elapsed time, map this estimate onto the corresponding reward probabilities, and select the action with the higher inferred reward probability. Together, the figure defines a class of inference problems in which reward is temporally structured but latent, requiring joint inference over time and reward probability rather than accumulation of sensory evidence.

A key unresolved question in this setting concerns the uncertainty in elapsed time estimation. Classic interval timing theories posit that uncertainty increases proportionally with elapsed duration, consistent with Weber’s law^4,22,32^, an assumption incorporated into influential models of anticipation^33^ and reward expectation^34^. In contrast, recent work demonstrates that task-relevant probability, rather than elapsed time per se, modulates temporal uncertainty^35,36^. This suggests an alternative hypothesis: temporal uncertainty may be shaped by expected reward, such that time points of higher reward probability are represented with greater precision than periods associated with lower reward. These alternatives make opposing predictions when reward probability varies systematically over time (Fig. 2a, b), enabling a direct model-based comparison.

**Fig. 2.**
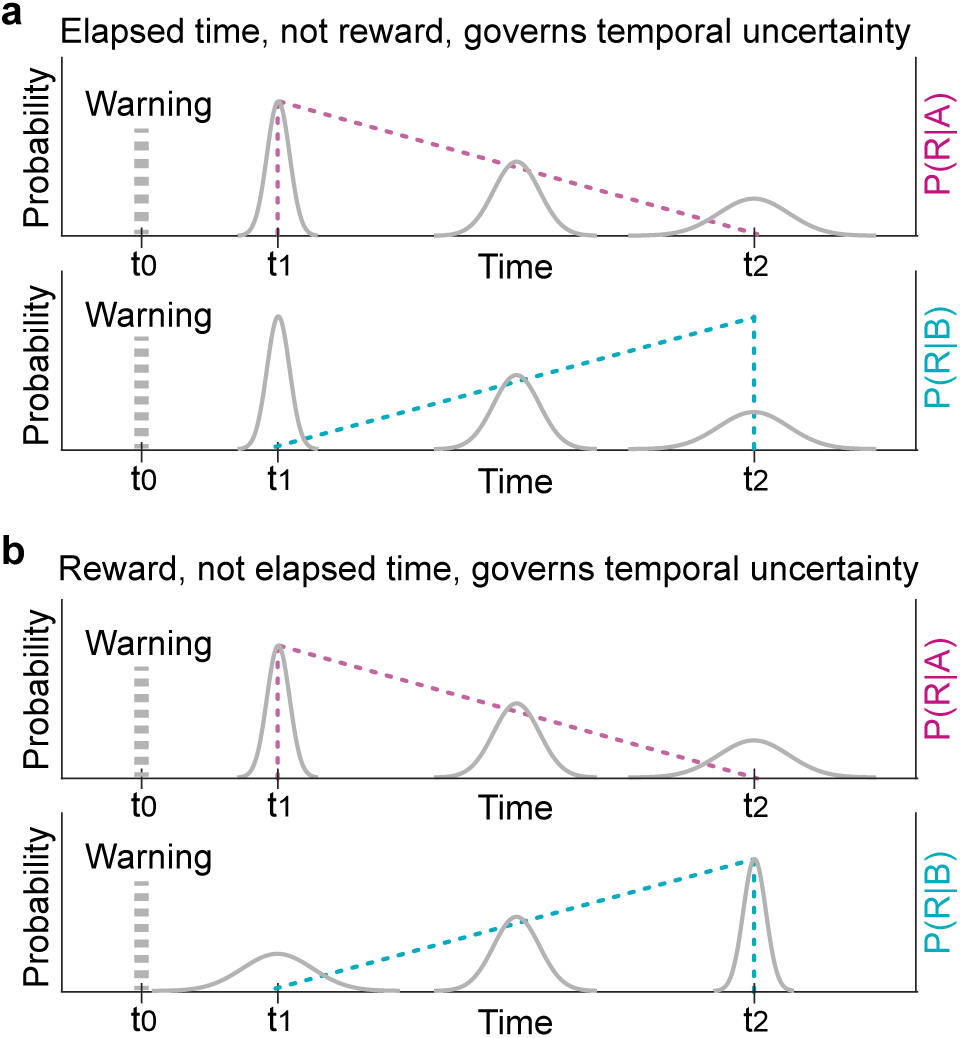
Competing hypotheses about temporal uncertainty. **a**, Temporal blurring (Weber-law scaling, Methods). Uncertainty in elapsed time estimation relative to a warning cue linearly increases with duration, independent of reward probability. Colored lines depict reward probability associated with actions A (top) and B (bottom). **b**, Probabilistic blurring (reward-contingent scaling, Methods). Uncertainty in elapsed time estimation is inversely related to reward probability, and does not depend directly on elapsed time itself, contradicting Weber’s Law. Under this hypothesis, temporal representations are more precise in periods associated with higher expected reward and less precise when reward probability is low. The two hypotheses make distinct predictions when reward probability increases over time (bottom panels), allowing model-based comparison of whether temporal precision is governed by elapsed time or by inferred reward structure.

A separate but related issue concerns how inferred probabilities are used to guide action. Extensive work has documented systematic deviations between objective probabilities and their subjective use in decision making^37-39^. These so-called probability distortions have been studied primarily in contexts where probabilities are explicitly stated (e.g. decision under risk^40^) and in static settings where probabilities are learned independently of time^41-43^. Whether similar transformations arise when reward probabilities must be dynamically inferred from temporal structure—rather than learned as stationary quantities—and how such transformations interact with temporal uncertainty, remains unknown.

Here we investigate behavior in a dynamic environment in which reward probabilities evolve over time and must be inferred from event timing. We formalize time-dependent choice as inference under dual uncertainty: uncertainty about elapsed time and uncertainty about reward as a function of time. Under the task structure, the reward-maximizing policy corresponds to a step-like transition at the objective crossover of reward probabilities (Fig. 1c), in words: "always choose the option associated with the higher reward probability". Implementing this policy requires temporally precise inference. Participants’ behavior approached optimality yet exhibited a systematic transformation of inferred reward probability, well captured by a linear mapping in log-odds space. Notably, this transformation operated on dynamically inferred, time-indexed probabilities rather than on explicitly stated or time-invariant probabilities, distinguishing the present framework from classic accounts of probability distortion. In parallel, temporal uncertainty was modulated by reward probability rather than by elapsed time, in contrast to Weber-law scaling.

Together, these findings identify two interacting computational principles. First, inferred reward probabilities are systematically transformed in the mapping to choice. Second, temporal precision is modulated by expected reward rather than elapsed time. By treating reward probability as a time-indexed latent variable and formalizing action selection as inference under dual uncertainty, the present framework links interval timing, reward estimation, and probability transformation within a unified account of time-contingent choice.

## Results

### Dynamic reward structure and behavioral performance

Participants performed a Set-Go task in which the time between two short visual cues (Go time) was drawn from a uniform distribution (Fig. 3a, Methods). In response to the Go cue, participants chose between two buttons and received immediate feedback, indicating whether their choice yielded a small monetary reward. Reward probabilities varied systematically over Go time (Fig. 3b): in all conditions, the probability of reward for a left choice decreased with time, whereas the probability of reward for a right choice increased. All conditions differed in their reward probability curves. This allowed us to investigate whether participants adapt to dynamic reward probabilities.

**Fig. 3.**
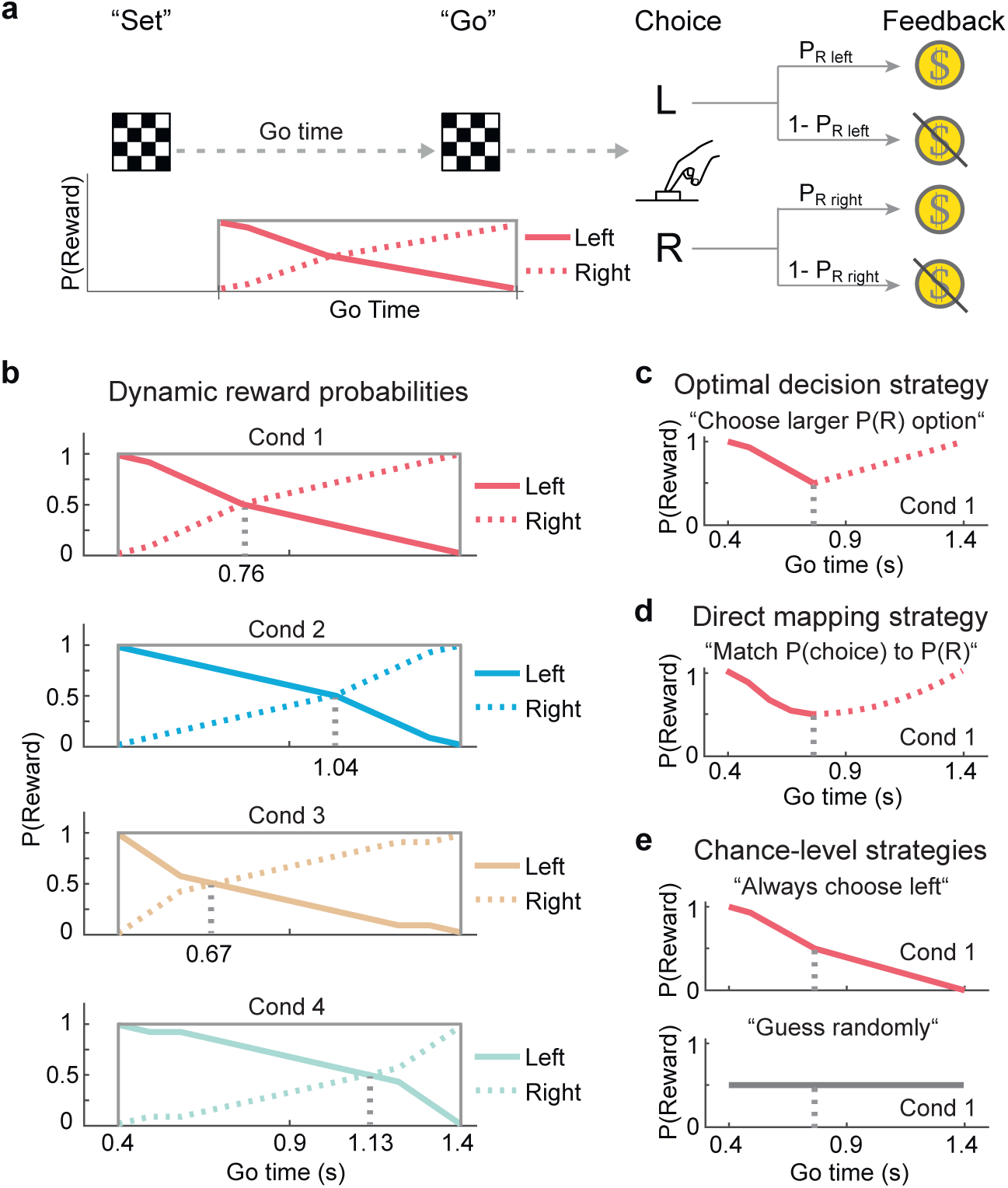
Time-based reward task and benchmark strategies. **a**, Set-Go task schematic. In each trial a Set cue was followed by a Go cue (50 ms checkerboards, Methods). The Set–Go interval (Go time) was drawn from a uniform distribution. Participants responded to the Go cue by choosing between either the left or right button and received feedback indicating reward outcome. **b**, Four dynamic reward conditions (separate blocks of trials). In all conditions, reward probability for a left choice decreased over Go time, whereas reward probability for a right choice increased. At each Go time, the two reward probabilities summed to 1. The dotted vertical line marks the crossover point where both left and right choices were equiprobably rewarded. **c**, Optimal strategy. At each Go-time point, the reward-maximizing policy is to choose the option with the higher reward probability, switching from left to right at the crossover time. Under perfect temporal knowledge, this deterministic policy yields an overall expected reward of P(R) = 0.78 (conditions 1–2) and P(R) = 0.77 (conditions 3–4). **d**, Mimicking strategy. If choice probability matches reward probability at each Go-time point (one-to-one mapping), expected reward is P(R) = 0.71 (conditions 1–2) and P(R) = 0.70 (conditions 3–4). **e**, Non-adaptive strategies. Always choosing one option or choosing randomly results in P(R) = 0.5.

At each Go-time point, the reward-maximizing policy is to choose the option associated with the larger reward probability (Fig. 3c). If this deterministic policy is strictly followed, this strategy yields an expected reward of *P(R)* = 0.78 (conditions 1 and 2) and *P(R)* = 0.77 (conditions 3 and 4). Because reward probabilities for left and right sum to one at each Go-time point, optimal performance requires inferring the reward function for at least one option as a function of elapsed time. Each condition features a specific Go-time point where *P*(*R*|*left*, *t*) = *P*(*R*|*right*, *t*) = 0.5, the crossover point (Fig. 3b, vertical dotted lines). The crossover point marks the point at which optimal choice switches from left to right, i.e. a step function that maximizes expected reward (Fig. 3c). Alternative strategies yield lower reward. Veridically matching choice probability to reward probability, e.g. for *P*(*R*|*left*, *t*) = 0.6, choose "left" in 6 out of 10 cases (mimicking), results expected reward of *P*(*R*) = 0.70–0.71 (Fig. 3d). Always choosing one option or responding randomly yields chance level performance (*P*(*R*) = 0.5; Fig. 3e).

Twelve participants completed the experiment, contributing 31,931 trials for analysis (Methods). Participants 1 to 6 performed conditions 1 and 2 and participants 7 to 12 performed conditions 3 and 4. Reward probability obtained from participants’ choices exceeded chance in all conditions (**Table 1**), ruling our stoic or random strategies. Thus, participants learned and exploited the dynamic reward structure.

**Table 1.**
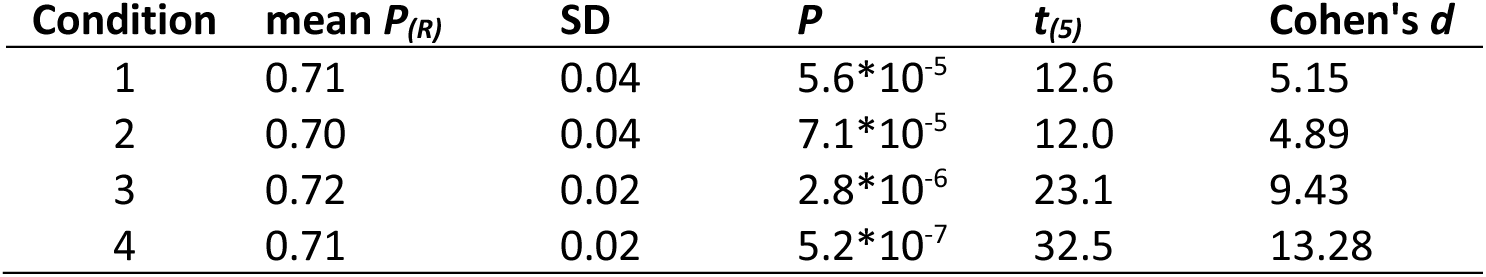
Participants’ mean reward probability significantly exceeded chance level *P(R)* = 0.5 (one-sided t test).

Across Go time, the temporal dynamics of reward probability obtained from participants’ choices approximated the optimal strategy, at both group-level (Fig. 4a) and single-participant level (**Fig. S1**). However, behavior did not fully implement the step-like policy. This raises the question whether participants’ choices followed the mimicking strategy (Fig. 3d). To investigate this hypothesis, we plotted the probability of left and right choices over Go time (Fig. 4b). Participants’ choice dynamics differ from the objective left and right reward probabilities in three ways: First, crossover points were shifted: participants overestimated the crossover time point in conditions 1 and 3 and underestimated it in conditions 2 and 4, effectively contracting subjective crossover toward the mean Go time. Second, choice probabilities deviated from a one-to-one mapping (mimicking) onto reward probability: before crossover, left choices were selected more frequently than implied by *P*(*R*|*left*, *t*) under the mimicking strategy, and less frequently thereafter (with the reverse pattern for right). Third, at the extrema of the Go-time range, large reward probabilities were slightly underexpressed in choice and small probabilities slightly overexpressed. Single-participant choice data closely reflect these group-level observations (**Fig. S2**).

**Fig. 4.**
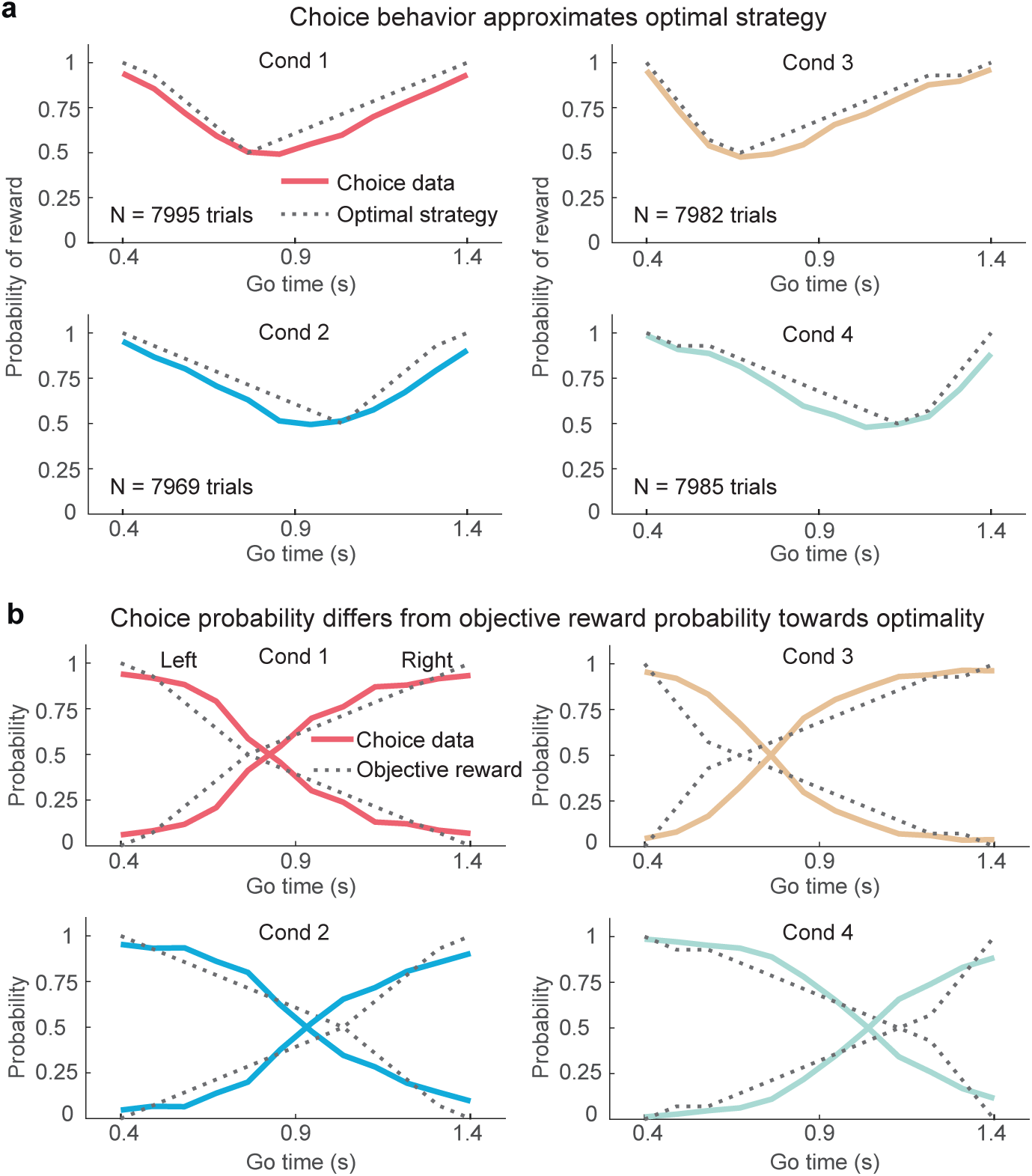
Decision dynamics across Go time. **a**, In all conditions, mean probability of reward over Go time, computed on participants’ choices (colored curves), approximates probability of reward according to the optimal decision strategy (dotted curves, **Fig. 3c**). Probability of reward was first computed on within-participant data, then averaged across participants. **b**, Mean choice probability over Go time computed individually for left and right choices systematically differs from objective reward probabilities towards optimality, i.e. towards a step function for left and right choices. Choice probability was first computed on within-participant data, then averaged across participants. Dotted curves represent objective reward probability over Go-time, as depicted in **Fig. 3b**.

Taken together, the analysis revealed that choice behavior does not mimic objective reward probability (Fig. 3d) and instead deviates towards optimal choice dynamics (step function). This raises the question how participants map the objective reward probability to their choices.

### Mapping reward probability to choice: a linear log odds transformation

To characterize how objective reward probability maps onto choice probability, we plotted the probability of a left choice against the objective reward probability given a left choice (Fig. 5). The resulting relationship was sigmoidal, inconsistent with both a step function (optimal policy) and a veridical one-to-one mapping (mimicking). Complementary plots for right choices showed the corresponding inverse relationship (**Fig. S3**).

**Fig. 5.**
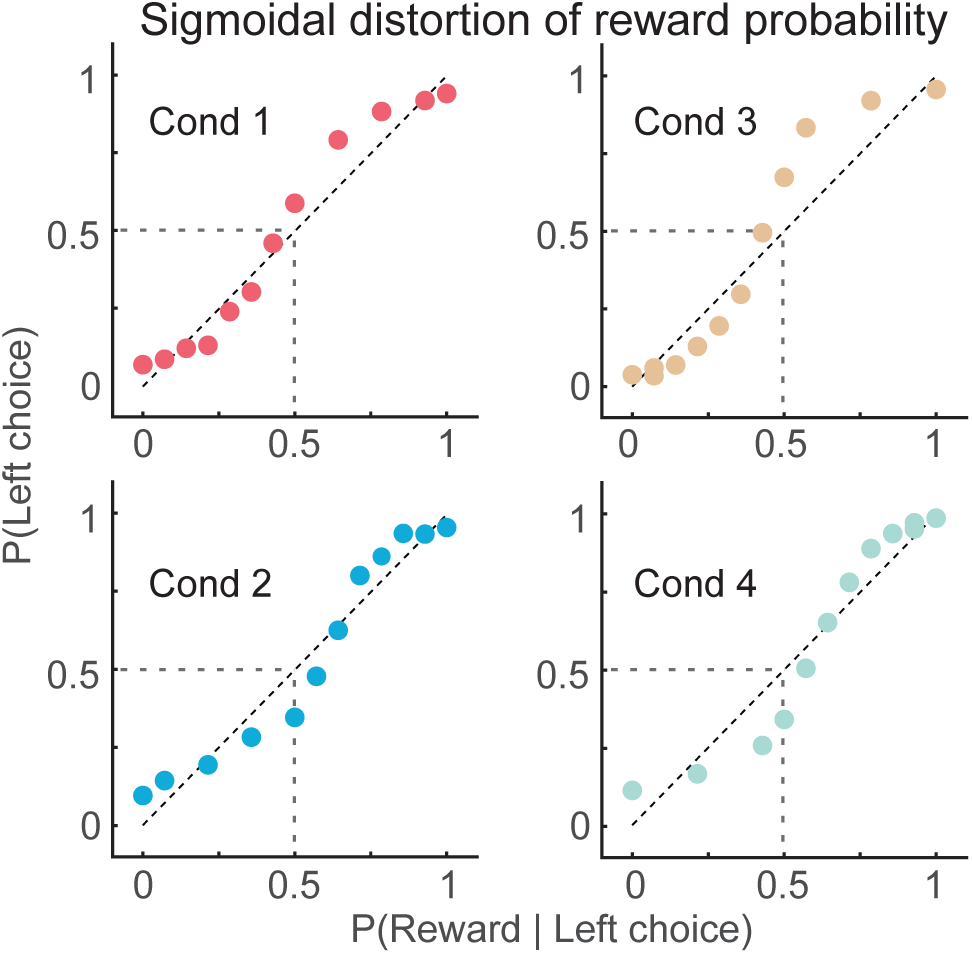
Sigmoidal distortion of objective reward probability. Plots of the mean probability of a left choice computed on participants’ data plotted against the probability of reward, given a left choice computed on stimuli, reveal a sigmoidal relationship between subjective (y-axis) and objective (x-axis) reward probabilities. Plots only show probabilities of participants’ left choices (first computed within-participant, then averaged across participants) and the corresponding left reward probabilities (computed on stimuli). The corresponding right choice probabilities are complementary (**Fig. S3**).

We modeled this mapping between objective reward probability over time and subjective choice probability using a log-odds linear operator (DLLO, Methods):

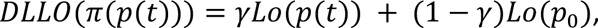

where *p(t)* denotes the objective reward probability at time t, *π*(*p*(*t*)) the corresponding choice probability, and 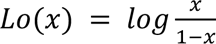 the log-odds^44^ transformation. The function DLLO defines a two-parameter familiy of transformations^45^, where *γ* is the slope parameter and *p_0_* is the fixed point of the transformation, the value of *p* that is mapped to itself. The subjective choice probability *π* is obtained by inverting the log-odds:

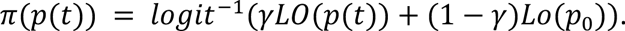

The function DLLO can capture both the one-to-one mapping of probabilities (mimicking) for *γ* = 1 and the optimal mapping for *γ* = *inf* (step function, Fig. 6a). For 1 ≤ *γ* ≤ *inf*, the mapping is sigmoidal as observed in the empirical pattern in Fig. 5 (n.b. for 0 < *γ* < 1, an inverted sigmoidal shape results, not observed in the data, limiting the sensible range to *γ* = [1, *inf*]).

**Fig. 6.**
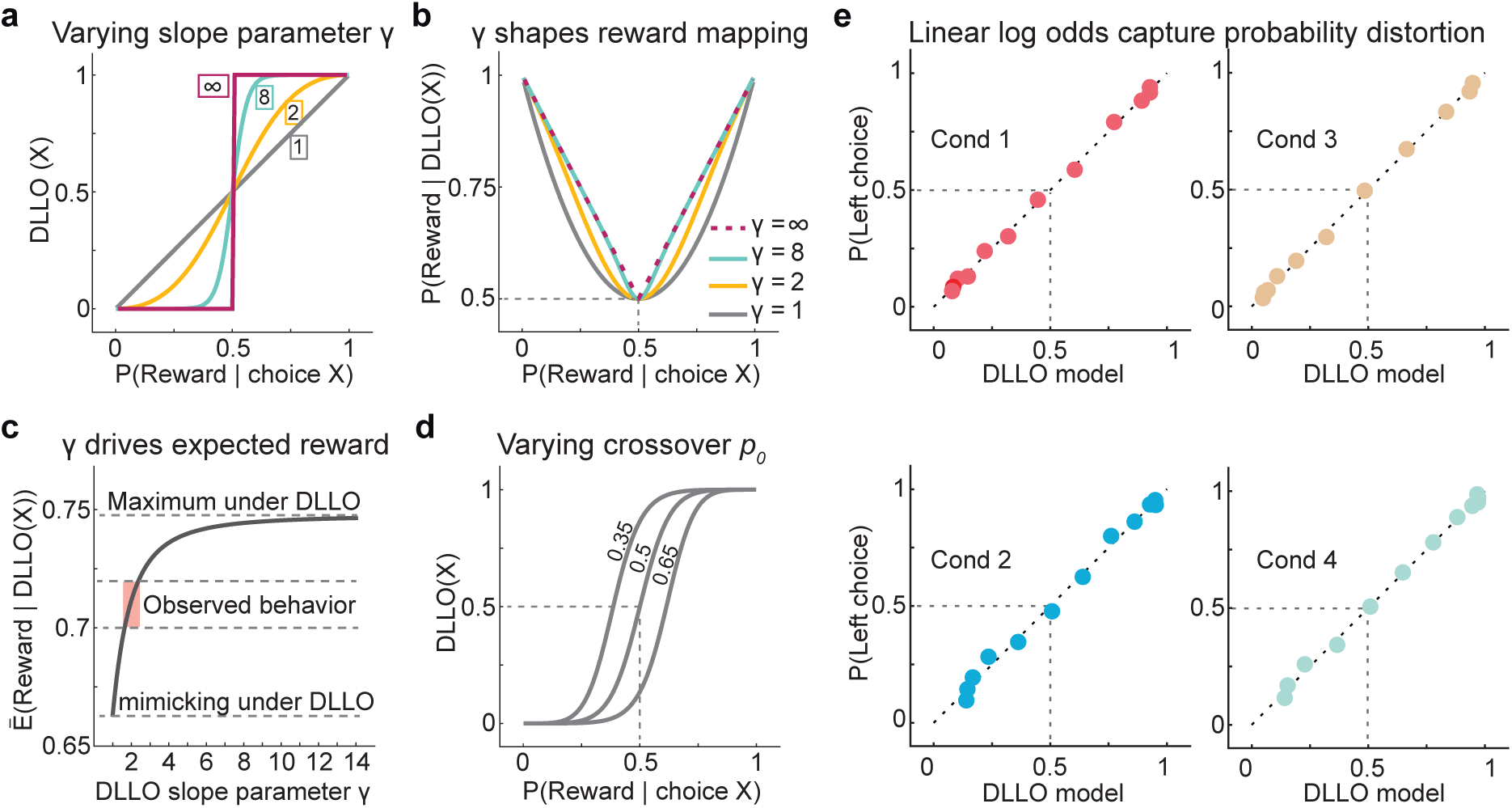
Linear log-odds mapping and its consequences for expected reward. **a**, DLLO transformation (Eq. 2) for slope parameters *γ* = [1, 2, 8, ∞] with crossover fixed at *p_0_* = 0.5. The function maps objective reward probability given choice X onto the probability of choosing X. At *γ* = 1, probabilities are mapped onto themselves (mimicking). As *γ* increases, the mapping becomes increasingly sigmoidal. In the limit *γ* → ∞ it approaches a step function, corresponding to a deterministic policy ("select the option with higher reward probability"). **b**, Effect of *γ* on the mapping between objective reward probability (x-axis), and subjective reward probability (y-axis) predicted by DLLO representation: as *γ* increases, reward probability increases. **c**, Expected reward as a function of *γ*. This panel provides a mechanistic illustration of the nonlinear relationship between *γ* and expected reward, computed under DLLO mapping across a uniform probability range (0.01–0.99) with *p_0_* = 0.5. Expected reward rises steeply for small increases in *γ* and asymptotically approaches the maximum attainable under this mapping (E(R) = 0.748). For *γ* = 1 (mimicking), expected reward is E(R) = 0.663. Dotted lines indicate the mimicking level, the asymptotic maximum under DLLO mapping, and the range of reward probabilities observed in participants’ behavior. The empirical values were obtained from task-specific reward distributions and estimated under the probabilistically blurred model; they are shown for qualitative comparison with the nonlinear *γ*–reward relationship illustrated here. Red vertical band indicates the range of *γ* values estimated from model fits (Table 2). **d**, DLLO transformation for different crossover parameters *p0* (*γ* fixed at 2) The parameter *p0* is the fixed point of the mapping, i.e. the reward probability mapped onto itself. **e**, Dynamic log-odds transformation of objective reward probability (fits of DLLO, x-axis) linearizes the relationship between objective reward probability and observed choice probability (y-axis, Methods).

**Table 2.** Parameter estimates from DLLO model fits to group-level choice probability.

| Condition | Choice | Adj. $R^2$ | $\gamma$ | $p_0$ |
| --- | --- | --- | --- | --- |
| 1 | left | 0.9979 | 1.77 | 0.46 |
|  | right | 0.9980 | 1.71 | 0.58 |
| 2 | left | 0.9927 | 1.75 | 0.73 |
|  | right | 0.9925 | 1.81 | 0.31 |
| 3 | left | 0.9987 | 2.20 | 0.44 |
|  | right | 0.9982 | 2.03 | 0.57 |
| 4 | left | 0.9971 | 1.84 | 0.66 |
|  | right | 0.9981 | 1.96 | 0.36 |

It is straightforward to compute the probability of reward that any DLLO mapping yields for a given choice X across the full range of reward probability *P_(R)_* = [0,1] (Fig. 6b). Note that for *γ* = *inf* the resultant curve for reward probability given DLLO(X) corresponds to the optimal decision strategy in Fig. 3c and for *γ* = 1 the curve corresponds to the "one-to-one" decision strategy (mimicking) in Fig. 3d. Numerically small values of e.g. *γ* = 2 yield curves that fall in between these two border conditions. As the *γ* numerically increases, the mapping quickly approaches the optimal strategy (see Fig. 6a and b for *γ* = 8). Thereby it becomes obvious that the expected reward probability, given DLLO mapping, is contingent on the slope parameter *γ* (Fig. 6c). Importantly, expected reward depends nonlinearly on *γ*: performance increases steeply for intermediate values (*γ* ≈ 1–3) and thereafter asymptotically approaches the optimal limit. Changes in *p_0_* result in a left/right shift of the mapping (Fig. 6d).

We fit DLLO models to group-level choice probabilities separately for left and right choices in each condition (Methods). Objective reward probabilities were first probabilistically blurred^36^ (Fig. 2b, Methods) to account for temporal uncertainty (see below). The DLLO-transformed values were then regressed onto observed choice probabilities. The model captured the data well, as demonstrated by the near-linear relationship between predicted and observed values (Fig. 6e).

Plotting the fit model to choice probability over Go time confirmed that DLLO captures adequately two key aspects of choice dynamics: (i) the model precisely fits participants’ subjective crossover point, (ii) the model captures choice dynamics over the entire range of Go time (Fig. 7). The model fits are supported by large values of adjusted R-squared (**Table 2**).

**Fig. 7.**
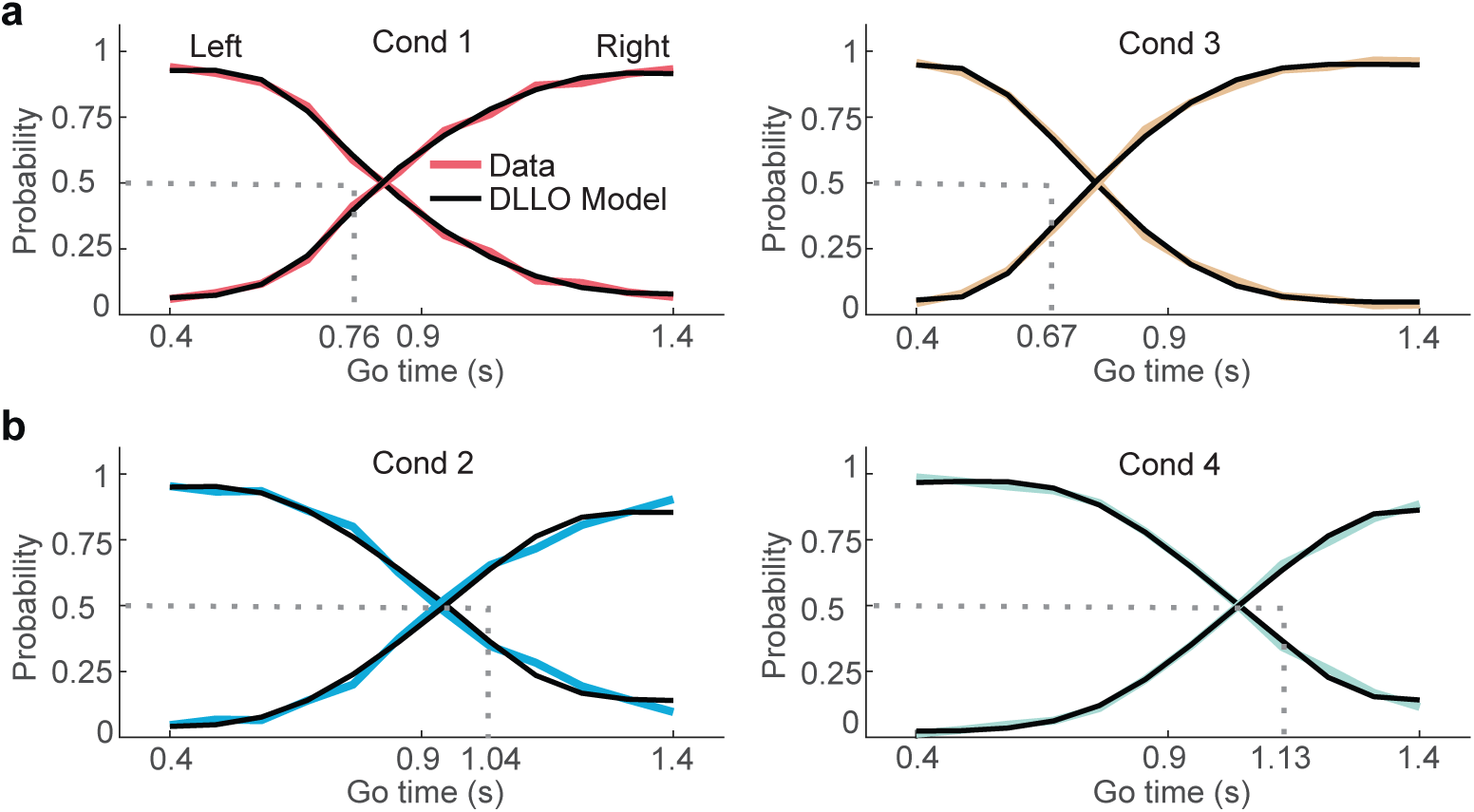
Linear odds of reward capture choice dynamics over time. **a**, In conditions 1 and 3, where participants over-estimated the crossover point (**Fig. 4b**, top row), fits of the DLLO model (Methods) capture participants’ choice dynamics over Go time. **b**, In conditions 2 and 4, where participants under-estimated the crossover point (**Fig. 4b**, bottom row), the DLLO model fits choice dynamics. Dotted lines depict objective crossover point between left and right reward probabilities as shown in **Fig. 3b**.

Across conditions, best-fitting *γ* values ranged from 1.71 to 2.20 (**Table 2**). It is in this range where the nonlinear relationship between *γ* and expected reward is most dynamic (Fig. 6c): Small increases in slope yield large increases in expected reward. Larger *γ* values produce diminishing returns, asymptotically approaching the optimal step-like policy that would require highly precise temporal estimates (Fig. 6a). Thus, behavior occupied a high-yield region of parameter space that captured most attainable reward without implementing an extreme step function.

In contrast, the crossover parameter *p_0_* varied across conditions but remained approximately symmetric around the objective crossover point of *P(R)* = 0.5 (**Table 2**). A simulation analysis revealed that these deviations had minimal impact on overall reward (**Supplementary Results**). Accordingly, *p_0_* likely reflects latent inference biases in time or reward representation rather than strategic tuning of the decision boundary (**Supplementary Note**). Together, these results identify *γ* as the principal adaptive degree of freedom governing how inferred reward probability is mapped onto choice.

### Robustness of DLLO fits at the individual participants’ level

To assess whether the observed transformations were driven by group averaging, we fit the DLLO model separately to each participant and condition. Fits were consistently strong across individuals (mean adj. R² = 0.98–0.99 across conditions, **Table S1**), indicating that the DLLO model captured choice probability at the single-participant level. The slope parameter *γ* exceeded 1 in all conditions (mean *γ* = 1.78–2.32, depending on condition), consistent with systematic distortion of inferred reward probability. Importantly, γ values exhibited moderate dispersion (SD = 0.32–0.87), yet remained within the high-yield regime identified in the reward analysis (Fig. 6c). No participant exhibited parameter estimates consistent with either a purely veridical mapping (*γ* ≈ 1) or the step policy (*γ* → ∞). The crossover parameter *p_0_* showed condition-dependent variation but with relatively modest spread (SD ≈ 0.10–0.16), replicating the group-level pattern. Full parameter estimates are reported in **Table S1**. These results demonstrate that dynamic distortion in log-odds space is a stable feature of behavior at both individual and group levels of analysis.

### Reward probability shapes temporal uncertainty

Because reward probability chances over time, inference requires a representation of elapsed time. Classical interval timing accounts assume that temporal variability scales with elapsed duration (Weber’s law)^4,32^, often implemented in modeling by a Gaussian blurring kernel whose standard deviation *σ* increases linearly with elapsed time, *σ* = *φ* · *t*, where *t* is elapsed time in seconds and *φ* is a constant, a Weber fraction for time estimation (Methods). We refer to this as *temporal blurring*.

Recently it was shown that in the temporal anticipation of sensory cues, uncertainty in time estimation is modulated by cue probability density^36^, rather than by elapsed time itself: Where cue probability is large, temporal estimates are precise and vice versa. This hypothesis is called *probabilistic blurring*. Here, probabilistic blurring is implemented by a Gaussian blurring kernel whose *σ* is inversely related to reward probability (Methods).

We compared both model variants: the temporally blurred DLLO model consistent with Weber scaling, and the probabilistically blurred model. The temporally blurred DLLO model under-estimates participants’ cross-over point (conditions 1, 2 and 4) and over- and under-estimates the probability of right choices over time in all conditions. (**Fig. S4**). In contrast, the probabilistically blurred DLLO model provided the best overall account of the data, capturing crossover shifts and full temporal dynamics as evidenced by **Figs. 6e** and Fig. 7 and **Tables 2** and **S1.** Analysis of residuals confirmed its superiority of the probabilistically blurred model (**Fig. S5**).

These results indicate that temporal precision in this task is better explained by reward-contingent scaling than by duration-dependent Weber scaling, suggesting that temporal uncertainty is structured by behavioral relevance rather than elapsed time alone.

## Discussion

This study identifies two interacting computational principles governing time-contingent choice when both reward probability and elapsed time must be inferred. First, inferred reward probabilities are systematically transformed in their mapping to choice, well captured by a linear transformation in log-odds space. Second, temporal uncertainty is modulated by reward probability rather than elapsed duration alone, inconsistent with Weber-law scaling. Together, these findings characterize behavior as joint inference over latent variables: elapsed time and reward probability over time. Both principles were robust at group and individual levels, indicating stable features of behavior.

### Dynamic distortion of inferred reward probability

Systematic distortions of objective probability are well established in static decision settings in which probabilities are explicitly stated or learned independently of time, as formalized in Prospect Theory and its cumulative extensions^37,38,41,46^. In these accounts, decision weights are nonlinear functions of known probabilities, typically overweighting small probabilities and underweighting moderate-to-large ones. Rank-dependent and dual-theory models formalized such effects as transformations of cumulative probabilities^46,47^. Parametric formulations characterized them using inverse-S-shaped weighting functions, most prominently the log-odds specification proposed by Prelec^38^, and related empirical demonstrations^48^. More recent work has shown that such distortions can arise from representational constraints and are well captured by linear transformations in log-odds space^49^.

The present results extend this framework in a critical respect. Here, reward probabilities are neither explicit nor stationary but must be inferred dynamically from event timing under temporal uncertainty. Probability transformation therefore operates on internally constructed, time-indexed representations rather than on on stated probabilities.

Behavior showed systematic overestimation of small inferred probabilities and underestimation of large ones, biasing choice toward—but not fully achieving—the optimal step-like policy. A linear log-odds mapping (DLLO) provided a compact and stable account across conditions (mean adj. R^2^ = 0.98–0.99). The slope parameter *γ* is the principal degree of freedom governing how inferred reward probabilities map onto choice. At *γ* = 1, behavior corresponds to veridical mapping (mimicking), whereas *γ* → ∞ approaches a deterministic step policy that implements the optimal switching rule. Participants consistently exhibited *γ* > 1, operating in a moderate range (*γ* ≈ 1.7–2.2).

Importantly, the consequences of this transformation can be quantified in expected reward. The *γ*–reward relationship was strongly nonlinear: modest increases from *γ* = 1 yield substantial gains in expected reward, whereas further increases produce diminishing returns, as performance asymptotically approached the optimal policy. Increases in *γ* primarily eliminate costly errors made at the extrema of the Go-time range, where reward probabilities are clearly high or low. Additional increases mainly affect choice near the crossover, where inferred probabilities are close to *P*(*R*) = 0.5 and gains are correspondingly small. Thus, behavior occupied a high-yield region of parameter space without requiring an extreme, discontinuous mapping. We caution that this interpretation is normative only with respect to the task’s reward dynamics.

### Crossover shifts and interpretation of *p_0_*

The optimal policy in our task is step-like: with perfect knowledge of reward probability over time, action should switch at the crossover where alternatives are equal in expected reward. However, because reward probabilities near this boundary are close to P(R) = 0.5, expected reward changes gradually around at the crossover point. Consequently, deviations in the subjective crossover location have limited effects on expected reward, reconciling the formal optimality of the step rule with diminishing returns in the *γ*–reward function.

The crossover parameter *p_0_* specifies the inferred reward at which choice is indifferent. Empirically, *p_0_* varied across conditions, shifting the subjective crossover relative to the objective *P*(*R*) = 0.5 boundary. Simulations showed that fixing *p_0_* = 0.5 while holding *γ* constant changed expected reward only minimally (ΔER < 2%), indicating that *p_0_* exerts limited influence on overall performance relative to *γ*.

Several non-exclusive factors may contribute to the observed shifts in *p_0_*: (i) Contraction to the temporal mean^50^, consistent with Bayesian timing accounts^6^ and scalar timing theory^4^. (ii) asymmetric reward gradients around the crossover (Fig. 3b), such that finite sensitivity (*γ* < ∞) displaces the subjective indifference point in reward-probability space. (iii) stable internal reference points in probability space that anchor choice independently of local reward gradients.

Our data do not permit a definitive separation of time-driven and reward-driven accounts. Because payoff differences near the crossover are small, these shifts incur limited cost. We therefore interpret *p_0_* primarily as reflecting latent inference biases rather than strategic reward tuning.

### Reward-contingent temporal uncertainty

Classic interval timing models posit that temporal uncertainty scales with elapsed duration^4,32^, implying that precision of temporal estimates increases as a function of time itself. In contrast, our results indicate that temporal precision is better explained by reward-dependent scaling: precision was highest during periods of high expected reward and lowest when reward probability was low. Thus, temporal precision was not a function of duration, but of the reward structure associated with time.

Previous work has emphasized that timing uncertainty constrains reward maximization, with agents taking normative account of their endogenous timing variabilty^51^. However, these account treats temporal uncertainty as a given constraint rather than as a quantity modulated by expected reward. A more recent article proposes that reward context can regulate the resolution of internal temporal representations via dopaminergic prediction-error signals: higher expected reward accelerates subjective time, whereas lower expected reward slows it^52^. This mechanism provides a principled route by which reward expectation could influence temporal resolution.

Our findings connect these perspectives by showing that reward expectation shapes temporal uncertainty directly at the behavioral level. In the DLLO model, probabilistic blurring scales inversely with reward probability. Although implemented using objective probabilities, this formalization approximates the observer’s learned reward–time structure rather than assuming explicit knowledge. Temporal precision is thus allocated according to inferred behavioral relevance, not elapsed duration alone.

Notably, reward-contingent temporal uncertainty and log-odds probability transformation operate at distinct but interacting computational levels. Temporal uncertainty constrains how precisely reward probabilities can be localized in time, whereas the log-odds mapping governs how those inferred probabilities guide action. Together, these coupled but conceptually separable constraints shape behavior when both time and reward are latent.

### Open questions and future directions

Several limitations point to directions for future work. First, we focused on binary choices with temporal-probabilistic structure. Extending this framework to richer action spaces^53^ or more complex stochastic temporal contingencies^35,54^ will be important. Second, our conclusions are based on behavior and neural mechanisms remain to be tested. A neural implementation of the log-odds computation appears plausible^45,55,56^, but how such transformations are neurally instantiated in time-contingent choice remains an open question.

More broadly, many real-world decisions require inferring reward structure from event timing. By characterizing how inferred probabilities are transformed and how temporal uncertainty is modulated in such settings, the present work delineates computational principles for decisions in which both time and reward must be internally reconstructed.

## Methods

The experiments were approved by the Ethics Council of the Max-Planck Society. Written informed consent was given by all participants prior to the experiment.

### Participants

12 healthy adults (5 female, 7 male), aged 20 to 33 years (mean age 22.6 years) completed the two-day experiment. All were right-handed and had normal or corrected-to-normal vision and reported no history of neurological disorder. Participants received € 14 per hour for participating, plus an additional bonus consisting of the cummulative monetary reward achieved in the experiment (see below).

### Task

In a visual Set-Go forced-choice task, a Set cue was followed by a Go cue. The time span between the onset of both cues, the "Go time", was randomly drawn from a uniform probability density function, spanning t = [0.4, 1.4] s. Participants were asked to respond as quickly as possible to the Go cue by choosing one of two buttons (left, right) using their left and right index fingers. Immediately after the button press, participants received visual reward feedback on their choices, indicating whether or not their choice is rewarded (green circle: rewarded, red circle: not rewarded). This choice feedback followed reward probability curves, one for left and one for right choices (see below).

### Visual stimuli

The Set cue consisted of two checkerboard patterns, which were presented simultaneously. One was positioned to the left of a central black fixation dot and the other on the opposite side. The Go cue consisted of two checkerboard patterns at the same location but with the black–white pattern reversed. Each checkerboard subtended 6.5 × 6.5° of visual angle and consisted of 7 × 7 black and white squares of equal size. The center of each checkerboard was positioned at a horizontal distance of 8.7° of visual angle and at a vertical distance of 0° from the center of the central fixation dot. Set and Go stimuli were each presented for 50 ms on a BenQ XL2420-B monitor (resolution 1,920 × 1,080, refresh rate 144 Hz), which was set to a gray background. All stimuli were generated using MatLab (the MathWorks) and the Psychophysics Toolbox (PTB-3)^57^ on a Fujitsu Celsius M730 computer running Windows 7 (64 bit).

### Experimental procedure

On each of two consecutive days, participants performed 8 experimental blocks, each consisting of 168 Set-Go trials. During each block, participants were instructed to foveate a central black fixation dot. Immediately after the button press, a small circle of green (choice rewarded) or red (choice not rewarded) color around the fixation dot was presented onscreen for 200 ms, providing visual reward feedback. The intertrial interval (ITI) was defined by the offset of the visual reward feedback and the Set cue of the following trial. The ITI was randomly drawn from a uniform distribution (range t = 1.4–2.4] s, discretized in steps of 200 ms). Participants were asked to restrict blinking to the ITI. At the end of each block, an onscreen number informed participants about the sum of rewarded trials they accumulated during the block. Participants were informed that each rewarded trial is monetized at the end of the experiment with a payment of 0.02€.

During the experiment, participants positioned their heads on a forehead-and-chin rest (Head Support Tower, SR Research Ltd.) at a fixed distance of approx. 60 cm from the computer monitor. An eye tracker (Eyelink DM-890, SR Research Ltd.) recorded participants’ eye movements at a sampling frequency of 500 Hz for fixation control. During the experiment, neural activity was recorded using an electroencephalography system (64-channel actiCAP with actiCHamp amplifier, Brain Products GmbH, Gilching, Germany, data not reported here).

### Go cue probability over time

The time between Set and Go cues, the Go time, was a random variable, drawn from a uniform distribution:

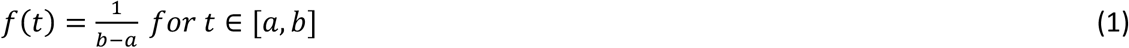

The uniform Go-time distribution was defined over the time span t = [0.4, 1.4] s at 12 equally-spaced individual Go-time points (Δt = 0.0909 s).

### Reward probability over time

At each Go-time point, two reward probabilities were defined for left and right choices. The left probability decreased over Go time, the "right" probability increased over Go time. The probabilities summed to one at each Go-time point. Four experimental conditions were constructed, each with a different set of "left" and "right" reward probability curves (Fig. 3b). Each condition featured a different crossover point, i.e. the Go time at which *P*_*left*_ = *P*_*ri*g*ht*_ = 0.5. Participants 1 to 6 (group 1) performed conditions 1 and 2, participants 7 to 12 (group 2) performed conditions 3 and 4. Each condition was presented for four consecutive experimental blocks before switching to the other condition. Half of the participants of group 1 began with condition 1, the other half with condition 2, and likewise for group 2.

### Data selection

Trials in which visual fixation was not maintained within a radius of 2.5° visual angle around the central fixation point for more than 0.3 s during the Go time were discarded for data analysis (n = 142 trials). Trials that featured an unreasonable long response time, i.e. RT > 1.45 s, were removed from the analysis (n = 183 trials). 31,931 trials remained for analysis.

### Modeling choice over time with a dynamic log-Odds linear Operator (DLLO)

Models of choice probability based on a linear transformation of the log-odds of reward (DLLO, Equation 2) were constructed to investigate choice dynamics over Go time. Three variants of DLLO predictors were considered: (i) non-blurred, (ii) temporally blurred, and (iii) probabilistically blurred.

### Non-blurred DLLO

The DLLO values were computed on the reward probabilities using Eq. (2):

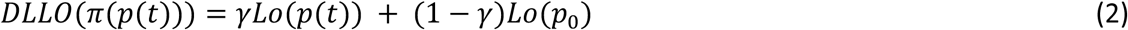

### Temporally blurred DLLO

Temporal uncertainty was modeled as a Gaussian convolution of the reward probability function *p*(*t*). The standard deviation of the Gaussian kernel scaled linearly with elapsed time:

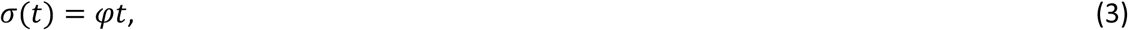

where *φ* is a scaling factor. The blurred reward probability at Go time *t*_*go*_ was computed as

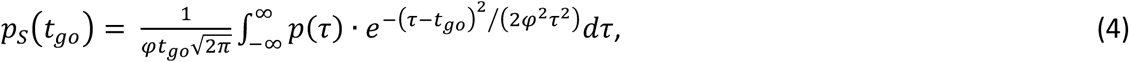

with cumulative form

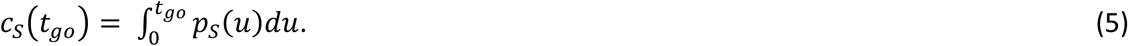

To ensure finite integration limits, *p*(*t*) was extended by three standard deviations at the shortest and longest Go times:

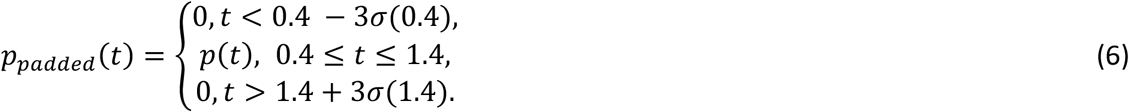

The scaling factor was fixed to *φ* = 0.11, corresponding the lower bound of values in the literature^58^, resulting in a comparatively small blurring effect. Reward probabilities were normalized such that 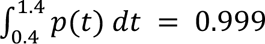 which reflects the certainty of a Go cue (no catch trials)^35^. DLLO values were computed from temporally blured reward probabilities using Eq. (2).

### Probabilistically blurred DLLO

Probabilistic blurring constitutes an alternative hypothesis to the temporal blurring described above^36^. In the probabilistic blurring model, uncertainty depended on reward probability rather than elapsed time: Go times with high reward probability are associated with low uncertainty in time estimation and vice versa, irrespective of the Go-time duration. The kernel variance was defined as a function of reward probability:

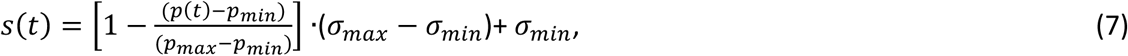

where *σ*_*min*_ = *φ* · 0.4 and *σ*_*max*_ = *φ* · 1.4. The value of *φ* was likewise set to *φ* = 0.11. The blurred reward probability was then computed as:

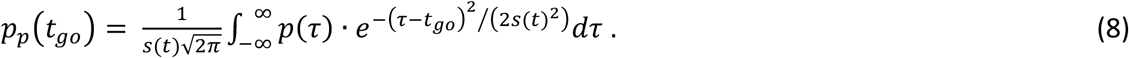

As in the temporal case, *p*(*t*) was extended by three kernel standard deviations beyond the [0.4, 1.4] s interval:

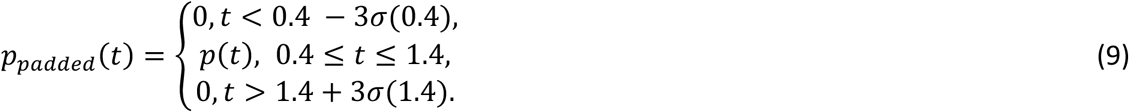

DLLO values were computed from probabilistically blurred reward probabilities using Eq. (2).

## Model fitting

For each reward probability curve, in all four conditions, the three to-be-fit variables (non-blurred, temporally blurred, probabilistically blurred DLLO) were computed for all parameter combinations *γ* ∈ [0.5, 2.5] in steps of 0.01 and *p*_0_ ∈ [0.1, 0.9] in steps of 0.01. Participants’ choice probabilities were computed within Go times within participants for single-participant fits and then averaged across participants for group-level fits. All variables (non-blurred DLLO, temporally blurred DLLO, and probabilistically blurred DLLO) were fit to choice probability data using a linear model. An Ordinary Least Squares (OLS) regression was employed for the computation of the regression coefficients using the MatLab (The MathWorks, Natick MA, USA) *fit* function. Model performance was quantified using adjusted R-squared.

## Acknowledgements

We thank Niels Hein for his help in data acquisition. We thank Georgios Michalareas for advice on the experiment and David Poeppel for advice and for project funding.

