## Supplementary File for "Dynamic distortion of inferred reward probability shapes choice over time"

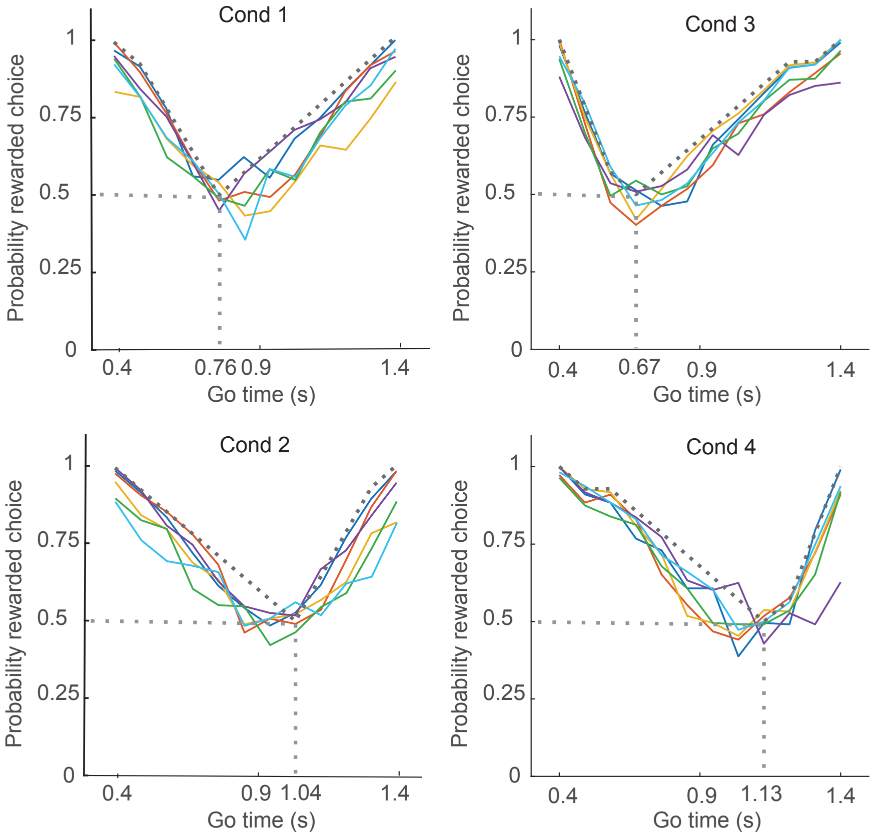


**Fig. S1 | Single-participant probability of rewarded choice over Go time.** In all conditions, mean reward probability over time, computed on within-participant choices (colored curves), approximates optimal reward probability (dotted curves). Vertical dotted line represents crossover point, i.e. the objective time point where left and right choices are rewarded equiprobably (P = 0.5, **Fig. 3b**).

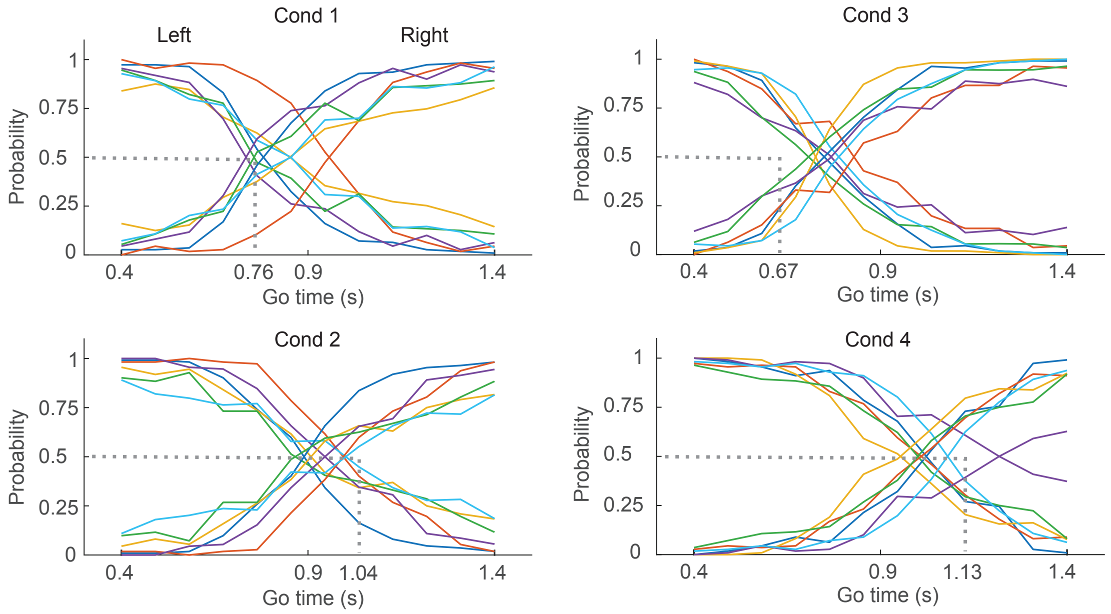


**Fig. S2 | Single-participant decision dynamics.** Mean choice probability over Go time computed individually for left and right choices (within-participant). Vertical dotted line represents crossover point of objective reward probabilities (**Fig. 3b**). Individual participants' choice dynamics are similar to group-level behavior (**Fig. 4b**). Most participants over-estimate the crossover point in conditions 1 and 3 and under-estimate it in conditions 2 and 4, i.e. contracting the crossover point towards the mean of the Go-time range.

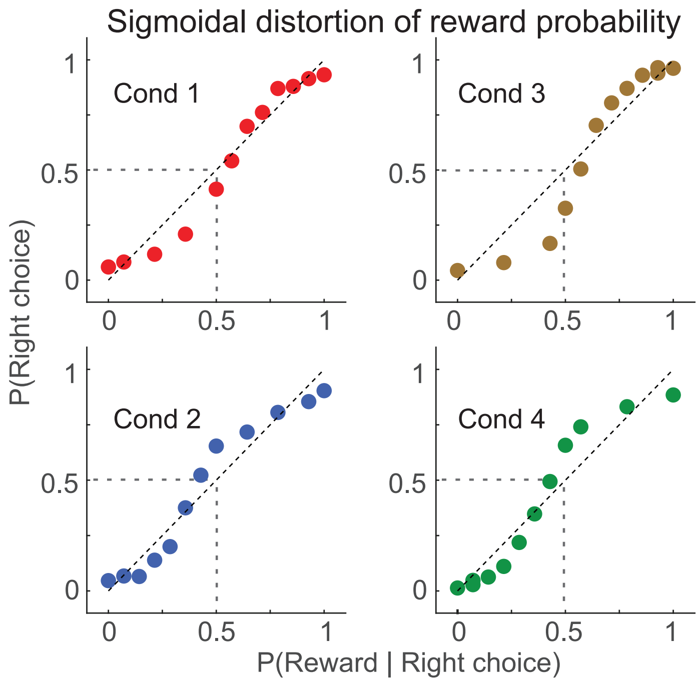


**Fig. S3 | Sigmoidal distortion of reward probability. a**) Plots of the mean probability of a right choice computed on participants' data plotted against the probability of reward, given a right choice computed on stimuli reveal a sigmoidal relationship between subjective (y-axis) and objective (x-axis) reward probabilities. Plots only show probabilities of participants' right choices (first computed within-participant, then averaged across participants) and the corresponding right reward probabilites (computed on stimuli).

**Supplementary Results**

**Quantifying the impact of $p_0$ on expected reward**

To assess the impact of the crossover parameter $p_0$ on expected reward (ER), we conducted a simulation-based analysis under the fitted DLLO model. The goal was to quantify how the condition- and choice-specific deviations of $p_0$ from 0.5 influenced reward outcomes. For each of the four experimental conditions, we used the best-fitting DLLO parameters estimated at the group level (**Table 2**). Specifically, we fixed the slope parameter $\gamma$ at its empirical value and used the corresponding fitted $p_0$ values to generate model-predicted choice probabilities over time. Expected reward was then computed at each Go-time point using the objective reward probabilities embedded in the task design:

$$ER_{fitted}(t) = P(choose\ left|t)\, p_L(t) + P(choose\ right|t) p_R(t),$$

where $p_L(t)$ and $p_R(t)$ denote the objective reward probabilities for left and right choices, respectively. We computed the expected reward contributions separately for left and right choices before summing to obtain total ER. Mean expected reward was obtained by averaging across time points within each condition.

To isolate the contribution of $p_0$, we repeated this procedure while holding $\gamma$ fixed at its fitted value but setting $p_0$ = 0.5 for all conditions. This manipulation effectively aligned the fixed point of the log-odds transformation with the objective crossover probability, thereby removing condition-specific shifts in the decision boundary while preserving the empirically observed parameter $\gamma$. The resulting change in mean expected reward ($\Delta ER = ER_{p0=0.5} - ER_{fitted}$) was small across all conditions: $\Delta ER = [0.0064, 0.0142, 0.0083, 0.0188]$. Thus, enforcing $p_0$ = 0.5 yielded only minimal improvements in expected reward, on the order of less than 1-2 percentage points. These results indicate that the observed condition-specific shifts in $p_0$ exert limited influence on overall performance relative to the effect of the slope parameter $\gamma$, which governs the steepness of the reward-to-choice mapping. Consistent with the reatively small numeric values of reward probability ($P \approx 0.5$) near the crossover point, moderate displacements of the decision boundary therefore incur only minor reward costs in the present task.

### Supplementary Note

#### Interpretation of the $p_0$ parameter

In the two-parameter log-odds model, $\gamma$ controls the sensitivity with which inferred reward probability is mapped onto action. In this context, sensitivity refers to how strongly small differences in inferred reward probability are amplified into differences in choice probability, effectively determining how steeply behavior transitions between actions as reward evidence changes over time. $p_0$ determines the centering of this mapping by specifying the inferred reward probability at which the observer is indifferent between actions. Unlike $\gamma$, which directly governs how sharply choice tracks reward probability and exhibits a strongly nonlinear relationship to expected reward, variations in $p_0$ have only a weak effect on reward in the present task. This is because $p_0$ primarily shifts the decision boundary along the time axis toward or away from the objective crossover point, where reward probabilities for the two actions are equal and marginal gains from increased precision are small. Consequently, the modest displacements of $p_0$ we observe in our modeling have a much smaller impact on expected reward relative to changes in $\gamma$.

$p_0$ varied systematically across conditions and choices (**Table 2**)., suggesting that it does not reflect tuning for reward maximization per se. Instead, $p_0$ is best understood as capturing biases in inference that may be orthogonal to reward optimization. These biases may arise from (i) contraction-to-the-mean effects in subjective time estimation[1,2], (ii) asymmetries in the reward–time gradients around the crossover point, (iii), or stable idiosyncratic reference points in inferred reward-probability space:

(i) Contraction-to-the-mean in subjective time estimation would instead bias internal time estimates toward the center of the experienced interval, producing a consistent displacement of inferred crossover points toward the midpoint of the Go-time range ($t = 0.9s$), even when objective reward structure varies across conditions.

(ii) Asymmetric reward gradients imply that equal temporal deviations on either side of the objective crossover point produce unequal changes in inferred reward probability. When choice sensitivity is finite ($\gamma < \infty$), this asymmetry can systematically displace the effective indifference point in reward-probability space, yielding condition-specific shifts in $p_0$.

(iii) Finally, idiosyncratic reference points may reflect stable internal criteria—such as preferred decision thresholds or priors over reward probability—that anchor choice behavior independently of task-specific reward gradients, leading to persistent offsets in $p_0$ with minimal impact on reward.

Importantly, because reward probability is close to P(R) = 0.5 near the crossover, such biases can persist without substantial cost to performance (**Supplementary Results**). Taken together, these considerations indicate that $\gamma$ constitutes the primary adaptive degree of freedom governing near-optimal performance under temporal uncertainty, whereas $p_0$ reflects secondary structure in inference or representation that has limited behavioral consequence in this task. Accordingly, we interpret systematic shifts in $p_0$ as informative about latent inference biases rather than as evidence for strategic tuning of the decision boundary.

**Table S1 | Parameter values of DLLO model fit to single-participant choice probability.**

| Condition | | Mean adj. $R^2$ | Std | Mean $\gamma$ | Std | Mean $p_0$ | Std |
|---|---|---|---|---|---|---|---|
| 1 | left | 0.9907 | 0.0085 | 2.0067 | 0.5301 | 0.4867 | 0.1515 |
| | right | 0.9905 | 0.0080 | 1.9483 | 0.4725 | 0.5467 | 0.1588 |
| 2 | left | 0.9826 | 0.0117 | 1.7833 | 0.3519 | 0.7317 | 0.1186 |
| | right | 0.9822 | 0.0124 | 1.8717 | 0.3634 | 0.3133 | 0.1171 |
| 3 | left | 0.9938 | 0.0062 | 2.3233 | 0.8662 | 0.4250 | 0.1078 |
| | right | 0.9930 | 0.0071 | 2.1467 | 0.8277 | 0.5833 | 0.1232 |
| 4 | left | 0.9896 | 0.0076 | 1.9250 | 0.3164 | 0.6600 | 0.1187 |
| | right | 0.9905 | 0.0072 | 2.0683 | 0.3190 | 0.3633 | 0.1046 |

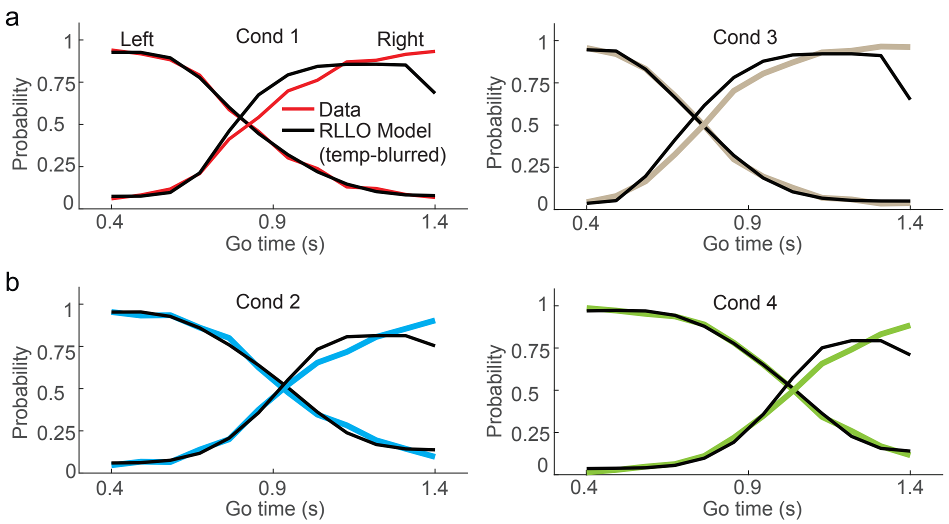


**Fig. S4 | Temporally blurred DLLO model systematically deviates from group-level choice probability across time. a**) In conditions 1 and 3, the temporally blurred DLLO model (**Methods**) under-estimates participants' cross-over point and over- and under-estimates the probability of right choices. **b**) In conditions 2 and 4, the temporally blurred DLLO model more accurately estimates participants' crossover point. The model over- and under-estimates the probability of right choices.

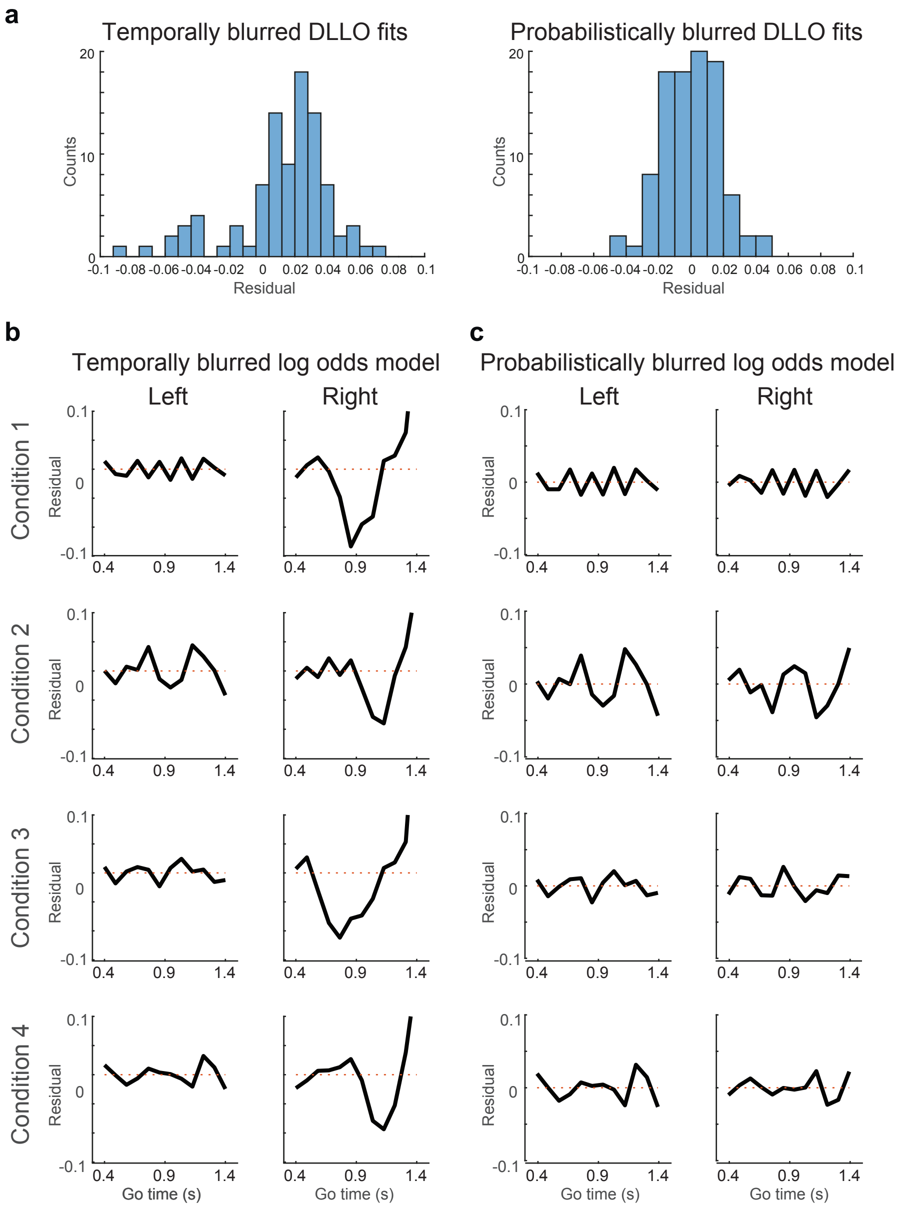


**Fig. S5 | Analysis of residuals from non-blurred, temporally blurred and probabilistically blurred DLLO models fit to group-level choice probability across Go time. a**) Distribution of model residuals differs between temporally blurred (left) and probabilistically blurred DLLO models. Temporally blurred DLLO model features numerically larger residuals than probabilistically blurred DLLO model. Distribution of residuals from temporally blurred DLLO model is skewed to the left. Residuals from probabilistically blurred DLLO model are close-to-symmetrically distributed around zero. **b**) Residuals from temporally blurred DLLO model. In the left-choice case, residuals do not feature an obvious pattern across conditions, reflecting the adequate model fit. In the right-choice case, there is an obvious trough and an increase in residuals towards the right extreme of the Go-time range, reflecting the systematic deviations between model and data (**Fig. S5**). **c**) Residuals from the probabilistically blurred DLLO model are numerically small across the range of Go times and do not feature systematic patterns across conditions, demonstrating an adequate model fit.

2 Vierordt, K. *Der Zeitsinn nach Versuchen*. (Laupp, 1868).
